# Computational target fishing by mining transcriptional data using a novel Siamese spectral-based graph convolutional network

**DOI:** 10.1101/2020.04.01.019166

**Authors:** Feisheng Zhong, Xiaolong Wu, Xutong Li, Dingyan Wang, Zunyun Fu, Xiaohong Liu, XiaoZhe Wan, Tianbiao Yang, Xiaomin Luo, Kaixian Chen, Hualiang Jiang, Mingyue Zheng

## Abstract

Computational target fishing aims to investigate the mechanism of action or the side effects of bioactive small molecules. Unfortunately, conventional ligand-based computational methods only explore a confined chemical space, and structure-based methods are limited by the availability of crystal structures. Moreover, these methods cannot describe cellular context-dependent effects and are thus not useful for exploring the targets of drugs in specific cells. To address these challenges, we propose a novel Siamese spectral-based graph convolutional network (SSGCN) model for inferring the protein targets of chemical compounds from gene transcriptional profiles. Although the gene signature of a compound perturbation only provides indirect clues of the interacting targets, the SSGCN model was successfully trained to learn from known compound-target pairs by uncovering the hidden correlations between compound perturbation profiles and gene knockdown profiles. Using a benchmark set, the model achieved impressive target inference results compared with previous methods such as Connectivity Map and ProTINA. More importantly, the powerful generalization ability of the model observed with the external LINCS phase II dataset suggests that the model is an efficient target fishing or repositioning tool for bioactive compounds.

## INTRODUCTION

Because most drugs exert their therapeutic effects by interacting with their *in vivo* targets, target prediction plays a pivotal role in early drug discovery and development, particularly during the era of polypharmacology (1). In the context of polypharmacology, the “magic bullet” (2) is likely an exceptional case, and *in silico* target prediction can be used to explore the whole therapeutic target space for a given molecule (3). This procedure might help deepen our understanding of the mechanisms of action, metabolism, adverse effects, and drug resistance of a molecule. By predicting targets of approved drugs, these clinically used chemicals can be repurposed for other diseases (4,5); for example, sildenafil (6) is used to treat erectile dysfunction but was first developed for the treatment of angina.

Targets of candidate molecules can either be identified via biochemical experiments, such as protein proteomic mass spectrometry (7), or predicted using computational approaches. Computational target prediction has gained momentum due to its low cost and high-throughput nature (8). The classical methods generally include ligand-based (9) and structure-based methods (10): the former methods mainly model drug-target interactions using features of small molecules, such as molecular fingerprints (11) and pharmacophores (12), and the latter methods often rely on molecular docking to unveil potential interactions between small molecules and proteins (13). Both of these methods rely on the similarity assumption: “similar molecules target similar proteins or vice versa” (14). However, this molecular similarity assumption does not always hold, e.g., structurally similar molecules can display different activities, such as the frequently observed activity cliffs (15). Moreover, ligand-based methods tend to exhibit decreased generalizability for new scaffold molecules that are not similar to any known drugs, and structure-based methods are limited by the lack of protein structures, inaccurate scoring functions, and a long computation time (16).

The rapid accumulation of transcriptional profiling data provides a new perspective for computational target prediction. For example, the LINCS L1000 dataset (17) is a comprehensive resource of gene expression changes observed in human cell lines perturbed with small molecules and genetic constructs. Several computational methods that involve the exploration of differential expression patterns have been proposed (18-27), and the strategies used in these methods mainly include comparative analysis, network-based analysis, and machine learning-based analysis (28). The comparative analysis-based methods infer targets based on gene signature similarities (17,24,26). An example is Connectivity Map (CMap), which assigns the target/MOA information of the most similar reference chemical/genetic perturbations to the new molecule by querying its gene expression signature against the reference L1000 library (17). The network-based approach systematically integrates gene expression profiles with cellular networks (19,20,29-32). For example, ProTINA applies a dynamic model to infer drug targets from differential gene expression profiles by creating a cell type-specific protein–gene regulatory network and provides improved prediction results compared with similar methods. Different machine learning algorithms have also been used in mining transcription profile data. Pabon et al. implemented a random forest (RF) model to explore the correlations between compound-induced signatures (CP-signatures) and gene knockdown (KD)-induced signatures (KD-signatures) from CMap and predict drug targets (33). Their study and that conducted by Liang et al. (34) revealed that the comparison of the differential expression patterns induced by chemical perturbation with those induced by genetic perturbation might shed light on potential information on the targets of a compound. Because these gene expression profile-based methods go beyond relying on the structural similarity between molecules, they are more suitable for discovering the targets of molecules with novel scaffolds. However, these studies generally focused only on differentially expressed genes and did not systematically consider the relationship among these differential genes in biological networks, the effects of compound concentrations and the cellular background, and differences in the time scales between compounds and shRNAs. Therefore, the major challenge in such investigations is that even if chemical and genetic perturbations interfere with the same target, the correlation between their gene signatures calculated using traditional methods might be very low because it is difficult to uncover the potential relevance of the gene signatures in biological networks under different conditions. To address this challenge, we propose a new graph convolution network (GCN) model, SSGCN. A trainable SSGCN was employed to integrate PPI information with raw signatures to derive graphical embeddings, and the results were then used to calculate the correlation between CP-signatures and KD-signatures. By concatenating the correlation results with the experimental CP time (the time from compound perturbation to measurement), dosages, cell lines, and KD time (time from KD perturbation to measurement), our model can predict drug targets across various cell lines, durations and dosages.

## MATERIALS AND METHODS

### 2.1 Data collection, preprocessing and splitting

#### 2.1.1 Data collection

The Library of Integrated Network-Based Cellular Signatures (LINCS) program, which is funded by the NIH, generates and catalogues the gene expression profiles of various cell lines exposed to a variety of perturbing agents in multiple experimental contexts. Both the LINCS phase I L1000 dataset (GSE92742) and the LINCS phase II L1000 dataset (GSE70138, updated until 2017-03-06) were downloaded from the Gene Expression Omnibus (GEO) (35,36) provided by the Broad Institute. These profiles were produced by a high-throughput gene expression assay called the L1000 assay, in which a set of 978 “landmark” genes from human cells was used to infer the expression values of the additional 11350 genes. This reduced “landmark” gene set enabled the LINCS program to generate a million-scale transcriptional profile. For the sake of connectivity analysis and convenience, our analysis focused on the level 5 signature data (replicate-collapsed z-score vectors) and used only real measured expression values to achieve further dimensionality reduction. The Python library cmapPy (37) was used to access the level 5 signatures from GCTx files.

STRING (38) is a database compiled for PPIs from both known experimental findings and predicted results. The human PPI network from the STRING v11 database was downloaded.

#### 2.1.2 Data preprocessing

The pipeline used for the preprocessing of the LINCS dataset is shown in Figure 1-a. (1) Profile signatures after perturbation with shRNAs (Phase I). shRNA experiments might exhibit off-target effects due to the “shared seed” sequence among shRNAs (17,39). To gain an abundant set of robust KD-signatures, we performed k-mean (k=1) clustering of the “trt_sh” signatures separated by the cell lines and KD time and maintained the core signature, which is the central signature of the cluster, as a representation of the corresponding cluster (25). The core signatures across eight data-rich cell lines (A375, A549, HA1E, HCC515, HT29, MCF7, PC3, and VCAP) were filtered to obtain the corresponding 978 “landmark” vectors. These 978 vectors constituted the input of curated KD-signatures. (2) Profile signatures after perturbation with compounds (phase I). The targets of the compounds were retrieved using the application programming interface (API) from the cloud platform (clue.io) provided by the Broad Institute. This retrieval resulted in 2032 compounds with 743 targets. Consistent with the curated KD-signatures, CP-signatures were curated by filtering “trt_cp” signatures out of the data-poor cell lines and non-landmark vectors. (3) Profile signatures after perturbation with compounds (phase II). We first filtered out those compounds contained in the phase I dataset and then retrieved the targets of the compounds from the aggregated ChEMBL bioactivity data on LINCS Data Portal through a representational state transfer API (40). The targets with pKd, pKi or pIC50 values greater than or equal to 6.5 were treated as the “true” targets (41). The retrieval resulted in 250 compounds with 488 targets. The raw signatures of these 250 compounds across eight data-rich cell lines (A375, A549, HA1E, HCC515, HT29, MCF7, PC3, and VCAP) were then extracted from the LINCS phase II dataset. As mentioned above, only the 978 “landmark” vectors were retained. We preferred to select the samples with a dosage of 10 μM and a duration of 24 h, and for the data without a dosage of 10 μM or a duration of 24 h, the gene signature for the closest conditions is used as an alternative.

**Fig. 1.**
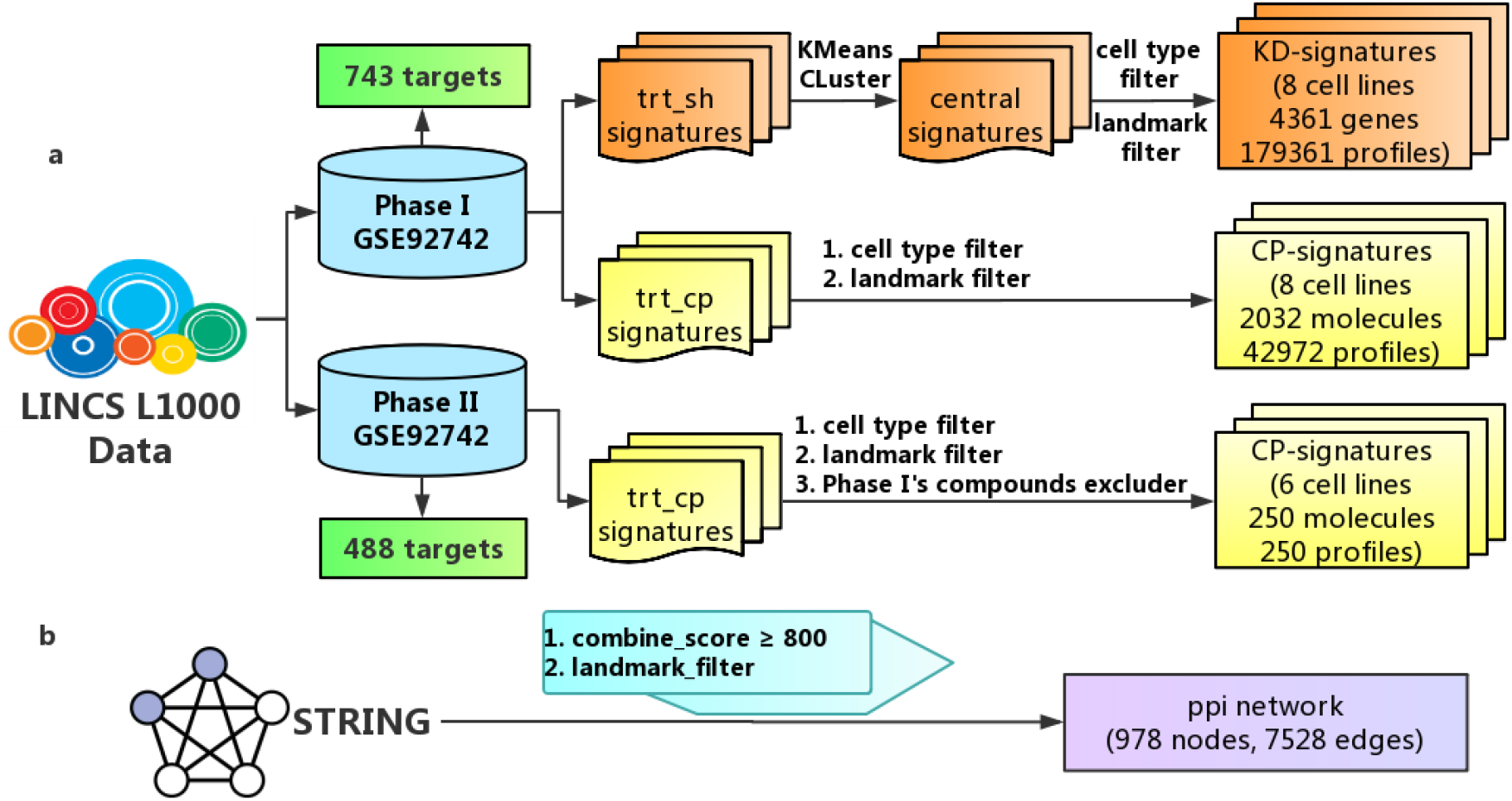
Pipeline of the data preprocessing. (a) Preprocessing pipeline for LINCS L1000 data. (b) Preprocessing pipeline for STRING PPI data.

We only kept the nodes present in the “landmark” gene set and the PPI edges with a “combined score” greater than or equal to 800. Accordingly, the curated PPI network consists of 978 nodes and 7528 edges (Figure 1-b).

#### 2.1.3 Data sampling

The test set compiled by Pabon et al., which contained 123 FDA-approved drugs that had been profiled in different LINCS cell lines and whose known targets were among the genes knocked down in the same cells, was used for benchmarking. Moreover, another benchmark dataset was prepared based on 250 compounds from LINCS phase II. The performance with these two external datasets can be used to evaluate our trained model in a real-world application scenario.

After excluding CP-signatures in these two external sets, we randomly sampled 80% of the remaining CP-signatures for training, 10% for validation and 10% for testing. To obtained a trained model close to real-world drug discovery settings, where inactive cases predominate, for each compound, three negative targets were generated for each positive target through a random cross combination of compounds and proteins.

### 2.2 Definition of the spectral-based GCN

An undirected graph G with 978 nodes was applied to represent the landmark PPI network. Each node in graph G represents a protein, and each edge represents a specific PPI interaction. Neighbourhood information is included in the edges. Traditional convolutional neural network structures are unfit for convolution operations on this graph, which is a non-Euclidian structure. Based on the Fourier transform of the graph and convolution theorem, spectral-based convolution operations on the graph can be applied to capture the properties of the graph network (42).

For a given graph G, its Laplacian matrix *L* can be defined as

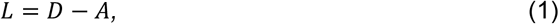

where *A* is the adjacency matrix of graph G and *D* is the degree matrix of graph G. In graph theory, the symmetric normalized Laplacian is more often used due to its mathematical symmetry. The symmetric normalized Laplacian *L_sys_* can be defined as

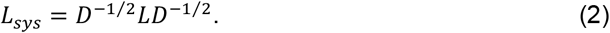

Based on the classical Fourier transform, we redefined the Fourier transform of the feature function in the node as the inner product of the function and the corresponding eigenvectors of the Laplacian matrix:

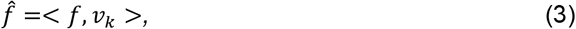

where *k* is the node on the graph, *f* is the feature function in node *k*, and *v_k_* is the eigenvector in the node of the Laplacian matrix. If spectral decomposition is performed on the Laplacian matrix, *L_sys_* can be expressed as

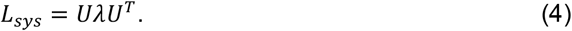

*U* is the orthogonal matrix of which the column vector is the eigenvector of the Laplacian matrix and *λ* is the diagonal matrix in which the diagonal is composed of the eigenvalues. The Fourier transform of the feature function *f* on the graph can then be rewritten as

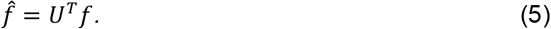

Because *U* is an orthogonal matrix, the inverse Fourier transform of function *f* on the graph can be written as

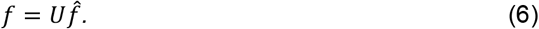

According to the convolution theorem in mathematics, a convolution procedure of two functions is the inverse Fourier transform of the product of their Fourier transforms. Defining *h* as the convolution kernel, the convolution operation on the graph can be expressed as

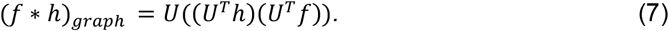

For the convolution operation in the first layer of the GCN, the Fourier transform of h is directly defined as the trainable diagonal matrix *ω*. Therefore, the convolution operation on the graph can be expressed as

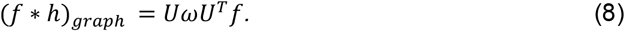

After the above derivation, the final form of the single layer of the spectral-based GCN can be expressed as

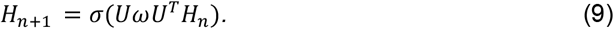

where *σ* is the activation function of the layer, *H_n_* is the input features of layer *n_th_*, and *H*_*n*+1_ is the output of layer (*n* + 1)_*th*_. According to the above definitions, the spectrum (eigenvalue) plays an important role in the convolution operation; thus, the GCN is called the spectral-based GCN. To effectively extract features and deeply learn from data, the multilayer perceptron can be connected to the graph convolution layer to increase the capacity of the model.

### 2.3 Model evaluation metric

The predictive performance of the model on the test set was evaluated using six classification metrics: accuracy, precision, recall, F1 score, area under the receiver operating characteristic (ROC), and area under the precision-recall curve (PRC). TP is the number of true positives, TN is the number of true negatives, FP is the number of false positives, and FN is the number of false negatives. All the metrics were calculated using the scikit-learn package (43), and a detailed introduction of the metrics is shown in Table 1.

**Table 1.** Introduction of the metrics.

| Metric | Description |
| --- | --- |
| Accuracy | $(TP+TN)/(TP+TN+FP+FN)$ |
| Precision | $TP/(TP+FP)$ |
| Recall | $TP/(TP+FN)$ |
| F1 score | $2 \times (\text{Recall} \times \text{Precision}) / (\text{Recall} + \text{Precision})$ |
| PRC | Area under the precision-recall curve |
| ROC | Area under the receiver operating characteristic |

## RESULTS

### 3.1 Spectral-based GCN for learning the network perturbation similarities

To capture the drug-target interactions and thus identify drug targets, we propose a SSGCN model that learns the undiscovered correlations between CP-signatures and the corresponding KD-signatures at the network level.

#### 3.1.1 Overall architecture of the model

Inspired by the study conducted by Pabon et al., the key idea of our target prediction model was to capture the correlations between chemical and genetic perturbation-induced gene expression in a more systematic manner. Based on this notion, targets of a compound can be predicted by comparing the corresponding perturbed gene expression profiles with a large number of KD-induced gene expression profiles that are publicly available. To learn potentially relevant information, as shown in Figure 2, two spectral-based GCNs were built: one for compound perturbation analyses, and one for gene perturbation analyses. This new architecture of the SSGCN model can also be divided into three main modules: the input module, the feature extraction module and the classification module. (1) The PPI network and differential gene expression profiles were the input of the first module. To unify information on the topology of the PPI network and the differential gene expression profiles, a property graph called a “gene signature graph” was constructed. Each node in the property graph represents a protein, and the property of each node was the corresponding differential gene expression value. Any two nodes are connected by an edge if two proteins can interact with each other. To represent compounds and targets, two gene signature graphs were constructed using compound a and gene perturbation data. (2) In the feature extraction module, the spectral-based GCN was used for graph embedding to integrate the PPI network topological structure information and differential gene expression profiles. Graph embedding provides a compressed representation of the gene signature graph. To obtain graph embeddings of the compounds and targets, two parallel GCNs were established for feature extraction. Because vector operations are more efficient than operations on graphs, after the gene signature graphs were transformed into graph embeddings, a simple linear regression layer could be used to characterize the degree of correlation between these two graph embeddings of compounds and targets. Gene expression profiles are also related to cell types, durations, and compound dosages (44). Therefore, correlation values terms of Pearson R^2^ concatenated with the experimental metadata (cell types, durations, and compound dosages) were fed into the classification module. (3) The classification module was composed of a fully connected hidden layer for extracting input features and an output layer for binary classification. The softmax function was applied in the output layer to compute the probabilities of whether the compounds show activity towards the potential targets (CPI scores). A label of 1 was assigned to a compound-protein pair if the compounds interacted with the corresponding protein, and a label of 0 was assigned to the opposite case.

**Fig. 2.**
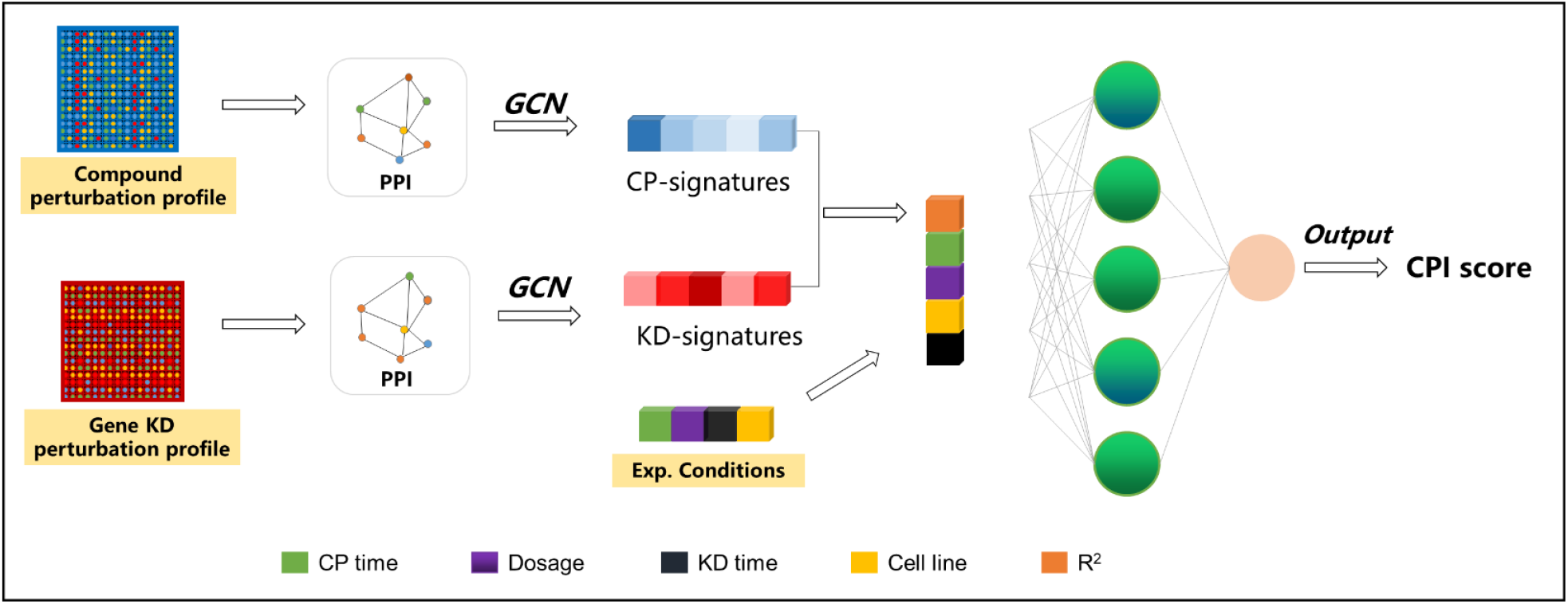
Architecture of the SSGCN.

The SSGCN model was implemented in the TensorFlow (45) framework (version TensorFlow-GPU 1.14.0) in Python 3.7.

#### 3.1.2 Training protocol

The SSGCN model was trained using the Adam optimizer for first-order gradient-based optimization. The model was trained to minimize the cross entropy between the label and the prediction result as follows:

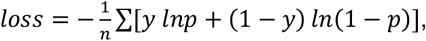

where *p* refers to the prediction result and *y* refers to the label. Early stopping (46) was used to terminate the training process if the performance of the model on the validation dataset shows no further improvement in specified successive steps, which helps selection of the best epoch and avoid overfitting.

#### 3.1.3 Target prediction with the SSGCN model

As shown in Figure 3, for a given compound **C**, the pipeline of predicting targets using the trained SSGCN model is as follows: (1) Obtain the compound perturbation gene differences on any of the eight cell lines by transcriptome profiling through ChIP sequencing (ChIP-Seq) and RNA sequencing (RNA-Seq) experiments or retrieving from public databases. (2) Extract a 978 CP-signature from the compound perturbation profile. (3) Feed the CP-signature and an existing KD-signature representing the gene perturbation profile of target **T** and their related experimental conditions, i.e., CP time, dosage, KD time, and cell line, to the trained SSGCN model for calculation of the CPI score of compound **C** and target **T**. (4) Repeat step 4 for the reference library of 179,361 KD-perturbation profiles. (4) Sort the potential targets by descending CPI scores. The top ranked targets are considered to be more likely to interact with compound **C**.

**Fig. 3.**
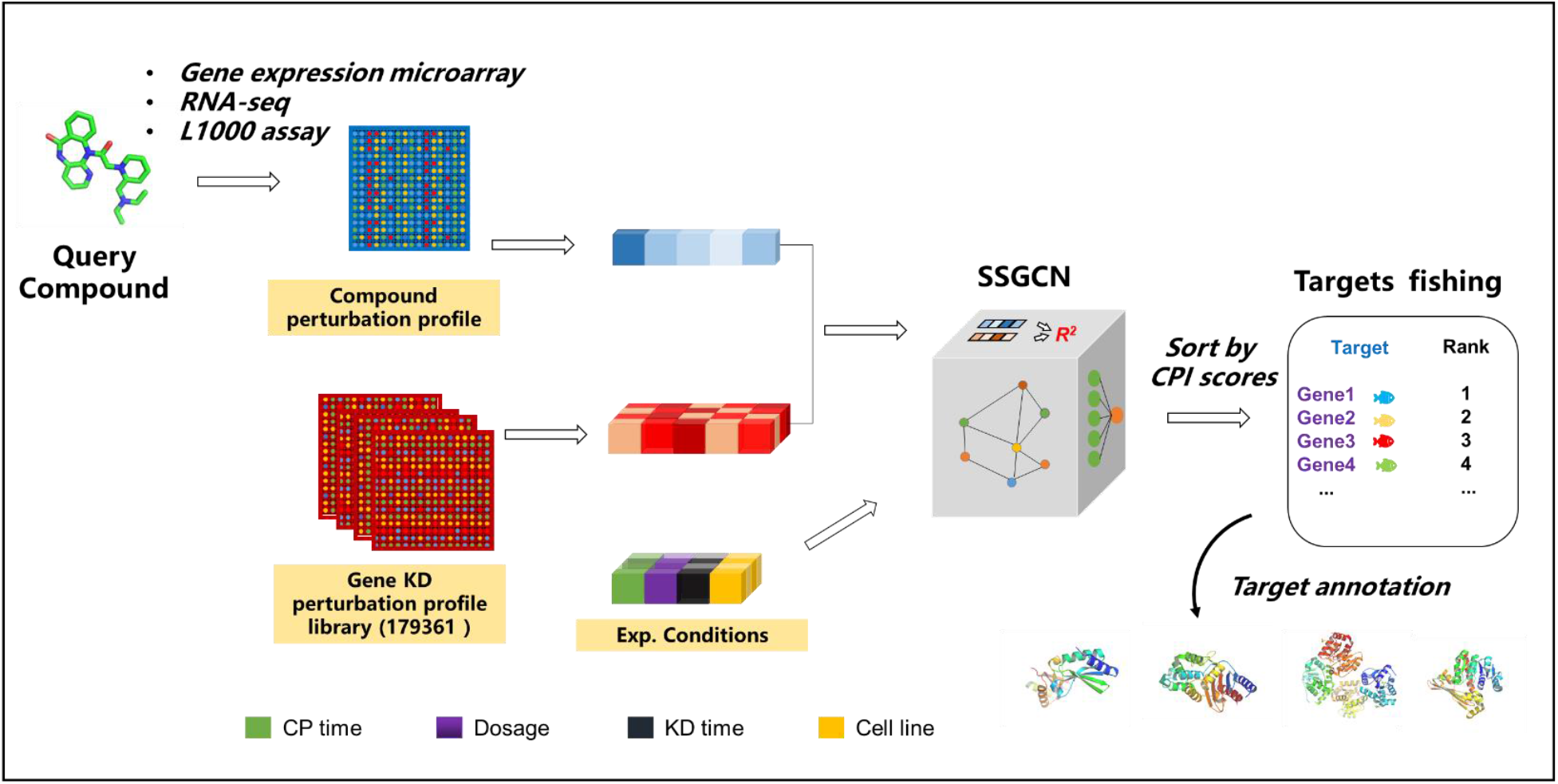
Pipeline of the target prediction

### 3.2 Optimization and internal test of the model

The SSGCN model is sensitive to the combination of hyperparameters. To optimize the model, as shown in Figure 4-a, different combinations of hyperparameters were evaluated with the validation dataset through grid searching. Because the number of negative samples was larger than that of positive samples (3:1), both PRC and F1-score are more suitable for evaluating the classification performance of the model. As summarized in Figure 4-a, the final model showed the best performance on the validation set with a learning rate of 10^−3^, a layer size of 2048 and a dropout (47) of 0.3. As shown in Figures 4-b and 4-c, the model yielded a PRC of 0.84 and an F1 score of 0.79 on the test dataset. Considering the high variance and noise in the gene signature data, the predictive performance of the model is impressive.

**Fig. 4.**
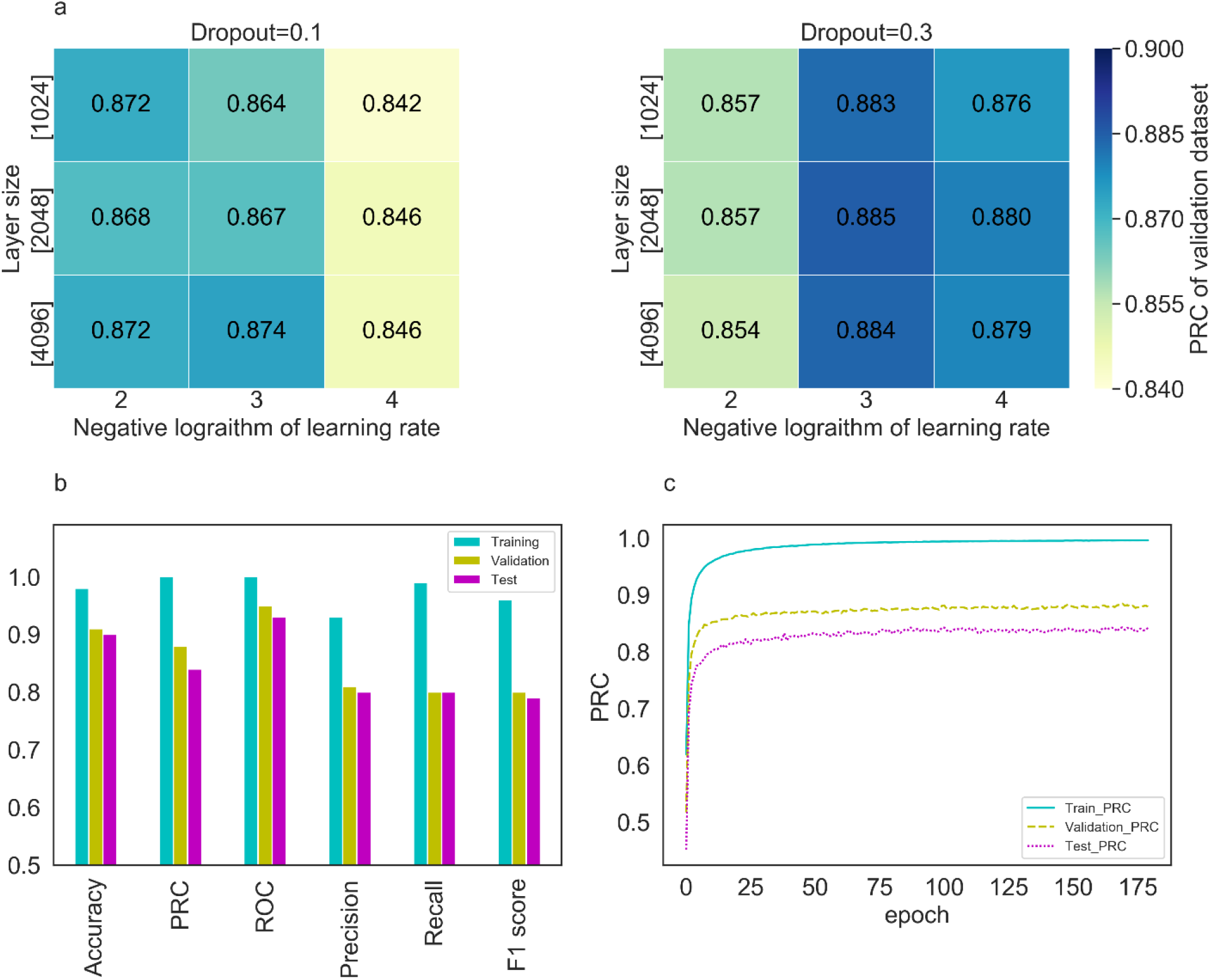
Model performance shown in (a) heat maps, (b) histogram graphs and (c) PRC curves.

### 3.3 External test and model comparison using LINCS phase I data

#### 3.3.1 Model performance and analysis using the external test set in LINCS phase I data

Although the model exhibited satisfactory results with the internal test dataset, we were more interested in its generalization ability for real-world target prediction tasks. Based on both the direct and indirect similarities between the chemical and KD perturbation signatures of cells, Pabon et al. applied an RF classification model to predict drug targets and constructed a dataset of 123 compounds and 79 targets, which could be considered a benchmark test for target prediction based on transcriptional profiles. To facilitate comparison, we used the same performance metric, Top N accuracy, to evaluate the performance of our model. This metric reflects the proportion of tested compounds whose any true targets can be correctly predicted among the top ranked N targets, and in this study, N values of 100 and 30 were evaluated. The prediction results reported by Pabon et al. were directly used for model comparison. For further comparison, a baseline model, CMap, was also implemented. For each compound in the external dataset, its top and bottom ranked 150 differentially expressed genes were used as the signature to query all the compounds in the LINCS phase I training data based on the CMap score. The value of the CMap score ranged from −100 to 100, where a large and positive value indicates that a reference compound could induce a signature similar to that induced by the query compound. Accordingly, all the known targets of the retrieved reference compounds with higher CMap scores were collected, and the top ranked 100 and 30 targets were assigned to the query compound as its candidate targets for calculating the top 100 and 30 accuracy values, respectively. Moreover, the network-based analytical method ProTINA was also benchmarked. Following the steps used in a previous study (19) and the provided code (https://github.com/CABSEL/ProTINA), the protein targets of the compound were ranked in descending order based on the magnitudes of the protein scores provided by ProTINA. It should be noted that different methods have different predicable target coverages. For SSGCN and the method reported by Pabon et al., the number of predicable targets corresponds to the number of different genes with available knockdown profiles in given cell lines. For CMap, the number of predicable targets is restricted to compound target-encoding genes. Among these methods, ProTINA covers more predicable targets because any genes with gene expression values can be considered potential targets.

To test our model, the gene expression profiles of these 123 compounds were excluded from the training dataset to avoid any potential information leakage. The remaining data were then used to train our model and predict targets for these 123 compounds according to the pipeline shown in Figure 3. As shown in Table 2, the top 100 accuracy values of the model in eight cell lines were higher than 0.7, and the model tested on the PC3 cell line showed the best prediction performance. Additionally, the prediction results obtained with SSGCN were better than those obtained with CMap and ProTINA. The relative ranks of the true targets were computed across eight cell lines. As shown in Figure 5-a, our prediction results on different cell lines were all significantly better than those reported by Pabon et al. (p<1e-10***).

**Table 2.** Target prediction performance on the external test set in 8 cell lines.

| **Methods** | **Number of compounds** | **Number of genes** | **Top 100 Accuracy** | **Top 30 accuracy** |
| --- | --- | --- | --- | --- |
| SSGCN (PC3) | 123 | 3980 | **0.84** | **0.71** |
| SSGCN (A549) | 123 | 3724 | 0.73 | 0.59 |
| SSGCN (MCF7) | 117 | 3688 | 0.82 | 0.64 |
| SSGCN (HT29) | 123 | 3665 | 0.72 | 0.46 |
| SSGCN (A375) | 122 | 3826 | 0.74 | 0.58 |
| SSGCN (HA1E) | 123 | 3801 | 0.80 | 0.63 |
| SSGCN (VCAP) | 120 | 4134 | 0.71 | 0.43 |
| SSGCN (HCC515) | 111 | 3522 | 0.77 | 0.63 |
| Pabon et al. | 123 | 3333 | 0.26 | 0.14 |
| CMap (PC3) | 123 | 2837 | 0.15 | 0.024 |
| ProTINA (PC3) | 120 | 10174 | 0.033 (0.058)* | 0.017 (0.033)* |

\* Because many more genes can be considered by ProTINA, the top 255 and 77 accuracy values, which denote the accuracy values at the same ratio of top 100 and 30 ranked targets, respectively, are also provided in parentheses for reference ( $255 = 100/3980 \times 10174$ , $77=30/3980 \times 10174$ ).

**Fig. 5.**
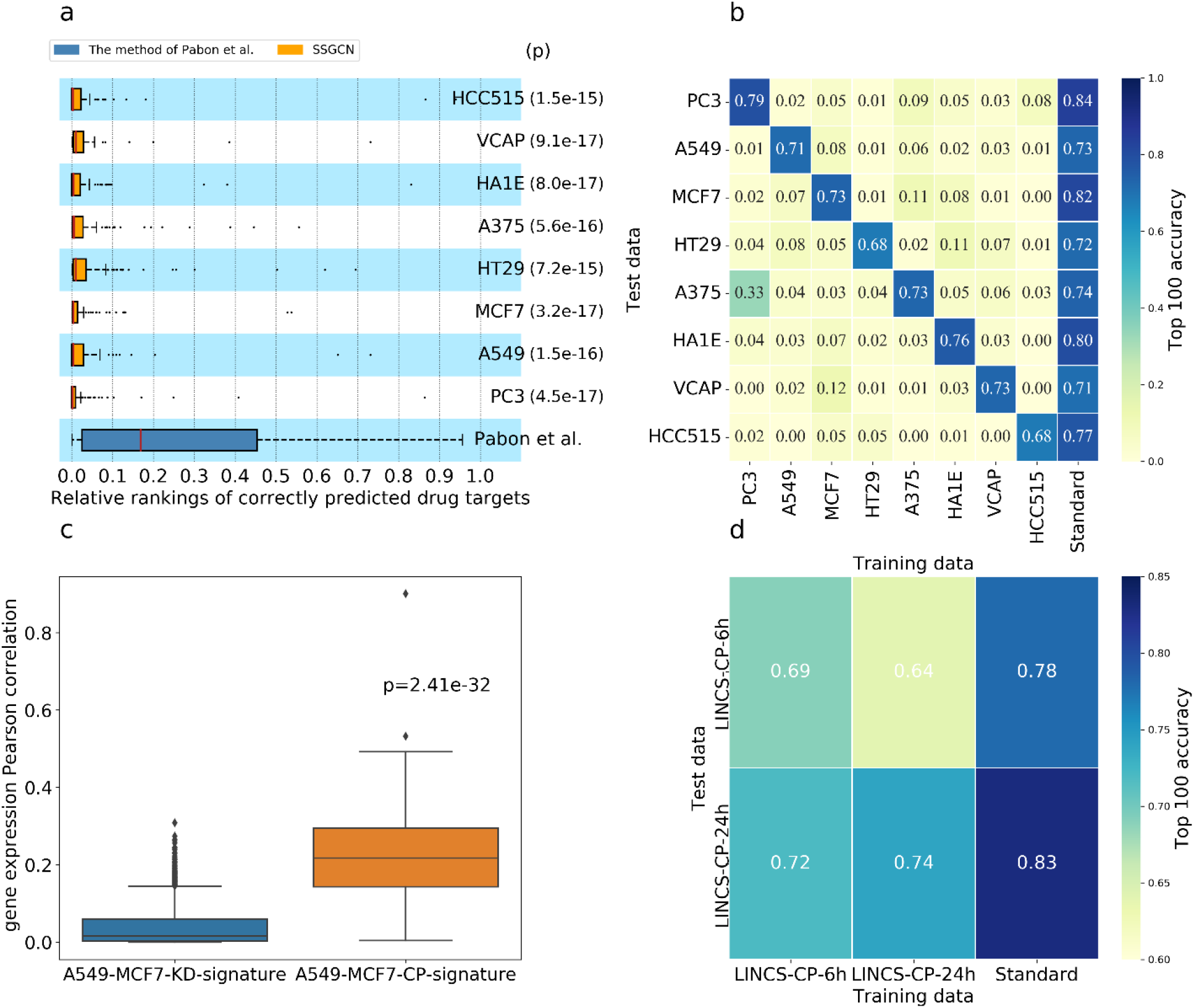
Model comparison and analysis. (a) Performance of the SSGCN models tested on different cell lines compared with that of the model developed by Pabon et al. (c) The correlation between the KD-signatures of A549 and MCF7 cells is significantly lower than that between the CP-signatures of these two cell lines. (d) Effects of the compound treatment time on target prediction performance.

To analyse the effects of the cell lines on the prediction performance, the datasets were split according to their cell lines (PC3, A549, MCF7, HT29, A375, HA1E, VCAP and HCC515). Eight individual submodels were constructed for each cell line and then separately tested on the external test dataset. As shown in Figure 5-b, these submodels could not make transferable predictions across cell lines, with the exception of the submodel trained with the transcriptional data of PC3, which showed only moderate prediction capability (Top 100 accuracy = 0.33) on A375. The limitation of these submodels can be attributed to the poor correlation between the KD-signatures among different cell lines when interfering with the same gene. As revealed in the original study (17), the similarity between shRNAs targeting the same gene is only slightly greater than random. Such similarity is even lower than that of signatures obtained after interfering with the same compound. Taking A549 and MCF7 as an example (Figure 5-c), the correlation of the KD-signatures between these two cell lines was significantly lower than that of the CP-signatures. In contrast, the standard SSGCN model trained with a combination of conditions showed excellent transferable prediction performance across all eight different cell lines, which makes it more useful in practical application scenarios. Similarly, to analyse the effects of the CP time on the target prediction, two individual submodels for different time scales (6 and 24 h) were built and tested. As shown in Figure 5-d, the models built from the LINCS-CP-6 h dataset achieved a Top 100 accuracy of 0.72 with the LINCS-CP-24 h test dataset, and those built from the LINCS-CP-24 h dataset achieved a Top 100 accuracy of 0.64 with the LINCS-CP-6 h test dataset. These results showed that the model could also make transferable predictions across CP times. In this study, the effects of the KD time on the target prediction were not analysed because most available KD-signatures were profiled at the same time (96 h, shown in Supplementary Table S1).

#### 3.3.2 The SSGCN model reveals a “deep correlation” between signatures

Using a computational normalization and scoring procedure, Pabon et al. demonstrated that drug targets can be predicted by comparing the correlations between chemical and genetic perturbation induced gene expression. Here, we further illustrated that target prediction can be significantly improved by encoding a PPI network and different experimental conditions when comparing these gene expression profiles. Therefore, it is of interest to investigate whether our SSGCN model could help reveal the “deep correlation” that cannot be revealed by conventional normalization and scoring. Intriguingly, the external test set contains gene expression profiles of 38 different NR3C1 antagonists and thus constitutes an ideal subset for comparing expression profiles after different chemical and genetic interferences on the same target. Using this subset, the target NR3C1 of 11 ligands was identified among the top 100 candidate targets by the method developed by Pabon et al. In comparison, for all these 38 ligands, NR3C1 can be successfully predicted within the top 100 targets by our SSGCN model. As shown in Figure 6, Raw R^2^ and KEGG Tanimoto coefficient represent two conventional correlation scoring methods for comparing gene expression values or KEGG pathway level features. No significant correlation was found between the chemical and shRNA-induced gene expression profiles using these two methods. In contrast, the correlations calculated by comparing graph embeddings from the PPI network and differential gene expression profiles, termed Deep R^2^, were markedly higher. These results highlight that our SSGCN model was able to determine the “deep correlation” between gene expression profiles upon heterogeneous drug treatments and explain why our model showed a markedly improved prediction performance in inferring targets based on transcriptional data.

**Fig. 6.**
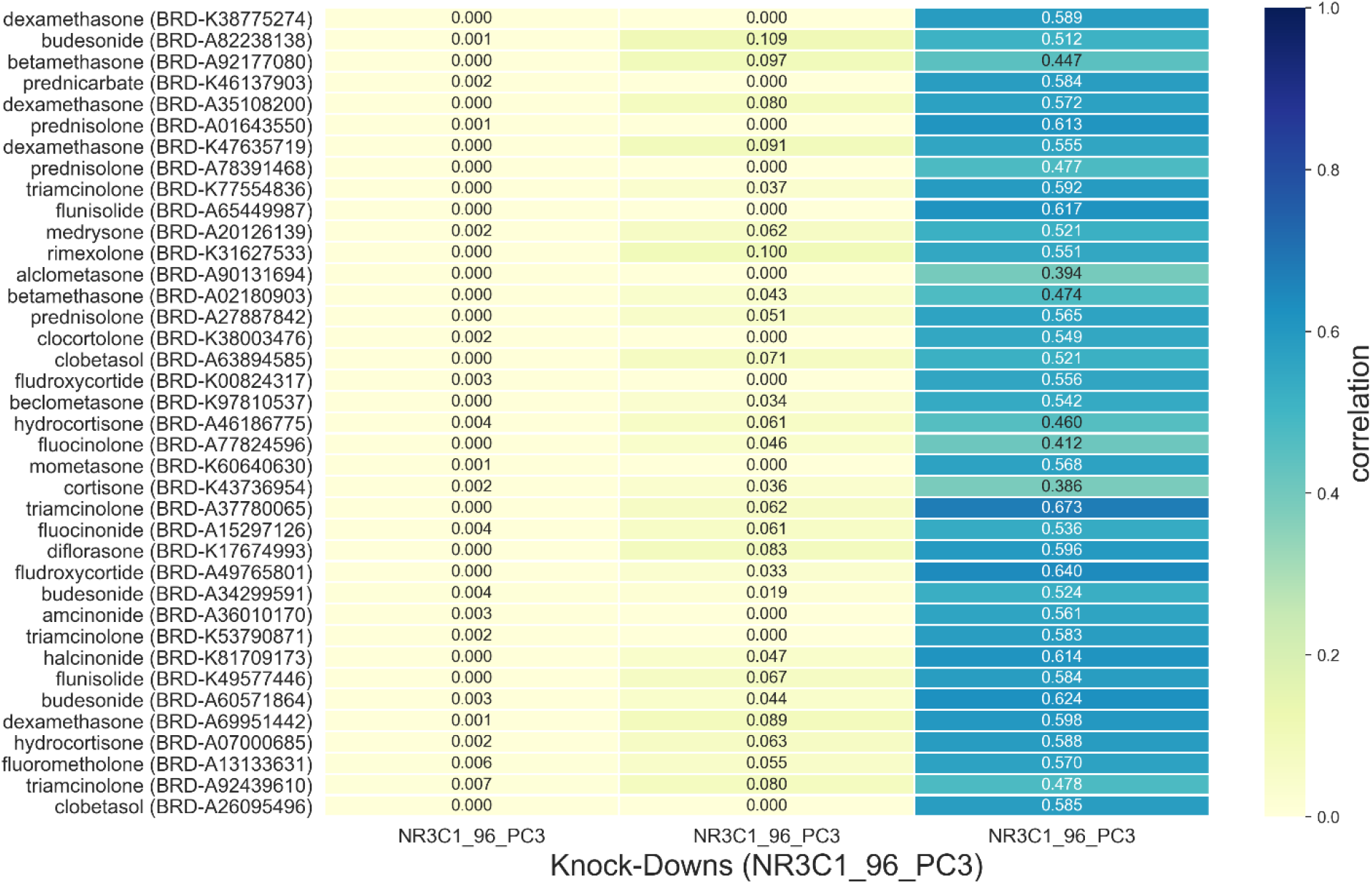
Correlation analysis of gene expression profiles. The RawR^2^, KEGG Tanimoto coefficient and deep R^2^ were used to represent the correlations of the raw gene expression values, KEGG pathway level features and graph embedding, respectively.

### 3.4 Model verification using LINCS phase II data

To further evaluate the generalization capability of the model in such a setting, as shown in Figure 1, LINCS phase 2 data were collected for stricter “time-split” testing (48). This dataset provides a more realistic prospective prediction setting in which the test data were generated later than the data used for modelling. After removing the overlapping compounds in the LINCS phase 1 data, the external test dataset includes 250 compounds and 488 targets. The trained model was employed to predict the targets of these compounds based on the target prediction pipeline shown in Figure 3. For comparison, a baseline model, CMap, was again implemented.

The time-split validation represents a more rigorous estimate of the model performance. As summarized in Table 3, the top 100 accuracy values of the SSGCN on the time-split external test set ranged from 0.51 to 0.66 in six cell lines. Although the accuracy declined slightly compared with the previous internal test with phase I data, which might be caused by different coverages of the target space (Supplementary Figure S1) and batch effects such as temperature, wetness and different laboratory technicians (17,49), the overall results of the SSGCN model are still highly impressive. In comparison, the baseline model using the CMap score for drug target prediction only yielded accuracy values lower than 0.31. We further performed a literature search for the new discovered targets of these external test compounds. For example, MAPK14 was ranked at the 26th position of the potential targets for saracatinib, and we searched European patents and found that the Kd value of saracatinib for MAPK14 is 0.332 μM. Similarly, MAPK1 was ranked at the 29th position among the potential targets of adenosine (50). This literature evidence further demonstrated the strong generalization capability of the SSGCN model for drug target prediction. For better visualization, a few external test compounds and their interaction network with the top 30 targets predicted by SSGCN are presented in Figure 7 (more details are provided in Supplementary Table S2). For example, the compound SB-939 is a potent pan-histone deacetylase (HDAC) inhibitor that inhibits class I, IIA, IIB and IV HDACs (HDAC1-11) (51). As shown in Figure 7-a, the top ranked 11 targets for this compound were all HDACs, which is accordance with the interacting targets reported previously. Alpelisib is an oral α-specific PI3K kinase inhibitor that has shown efficacy in targeting PIK3CA-mutated cancer (52), and its combination with fulvestrant has recently been approved by the US Food and Drug Administration for the treatment of metastatic or otherwise advanced breast cancer. Interestingly, the top ranked 30 targets of alpelisib are all types of different kinases, and PIK3CA can be successfully identified among the top three candidates. As a selective bromodomain-containing protein (BET) inhibitor, PFI-1 reportedly interacts with BRD4 with an IC50 of 0.22 μM (53). As shown in Figure 7-c, BRD4 was ranked third in the list of candidate targets. Moreover, our model predicted that PFI-1 might show cross-activity with a range of kinases. Because an increasing number of studies have shown that BRD4/BET inhibitors and kinase inhibitors might act synergistically in a range of cancer types (54), the predicted off-target interactions with kinases might provide clues and starting points for further study of related dual functional inhibitors (55-58). In some cases, the predictions were unsuccessful, e.g., ATM and RAD3-related (ATR) kinase if a reported target of VE-821, but this target was ranked at the 1594th position. As shown in Figure 7-d, the top 30 ranked targets identified by SSGCN cover a wide range of protein categories, including kinases, GPCRs and ion channels. Because compounds with smaller molecular weights might show promiscuity across different target families, we cannot rule out the possibility that VE-821 interacts with the predicted targets, but none of these interactions are supported by reported experimental evidence. This example also suggested that the candidate target list should be refined through further experimental verification and combination with other complementary methods, such as structure-based or similarity-based approaches. Overall, as indicated in Table 3 and Figure 7, it can be concluded that the SSGCN model shows strong generalization ability for inferring targets of previously unevaluated compounds and provides insights on cell-level transcriptomic responses to chemical intervention and related polypharmacological effects.

**Table 3.** Target prediction performance on the LINCS phase II data in 6 cell lines.

| **Cell lines** | **Number of**<br>**compounds** | **Number of**<br>**genes** | **Top 100**<br>**accuracy**<br>**(SSGCN)** | **Top 100**<br>**accuracy**<br>**(CMap)** |
| --- | --- | --- | --- | --- |
| PC3 | 249 | 3980 | 0.53 | 0.29 |
| A549 | 41 | 3724 | 0.66 | 0.31 |
| MCF7 | 240 | 3688 | 0.53 | 0.24 |
| A375 | 245 | 3826 | 0.51 | 0.30 |
| HA1E | 238 | 3801 | 0.56 | 0.27 |
| HCC515 | 39 | 3522 | 0.65 | 0.12 |

**Fig. 7.**
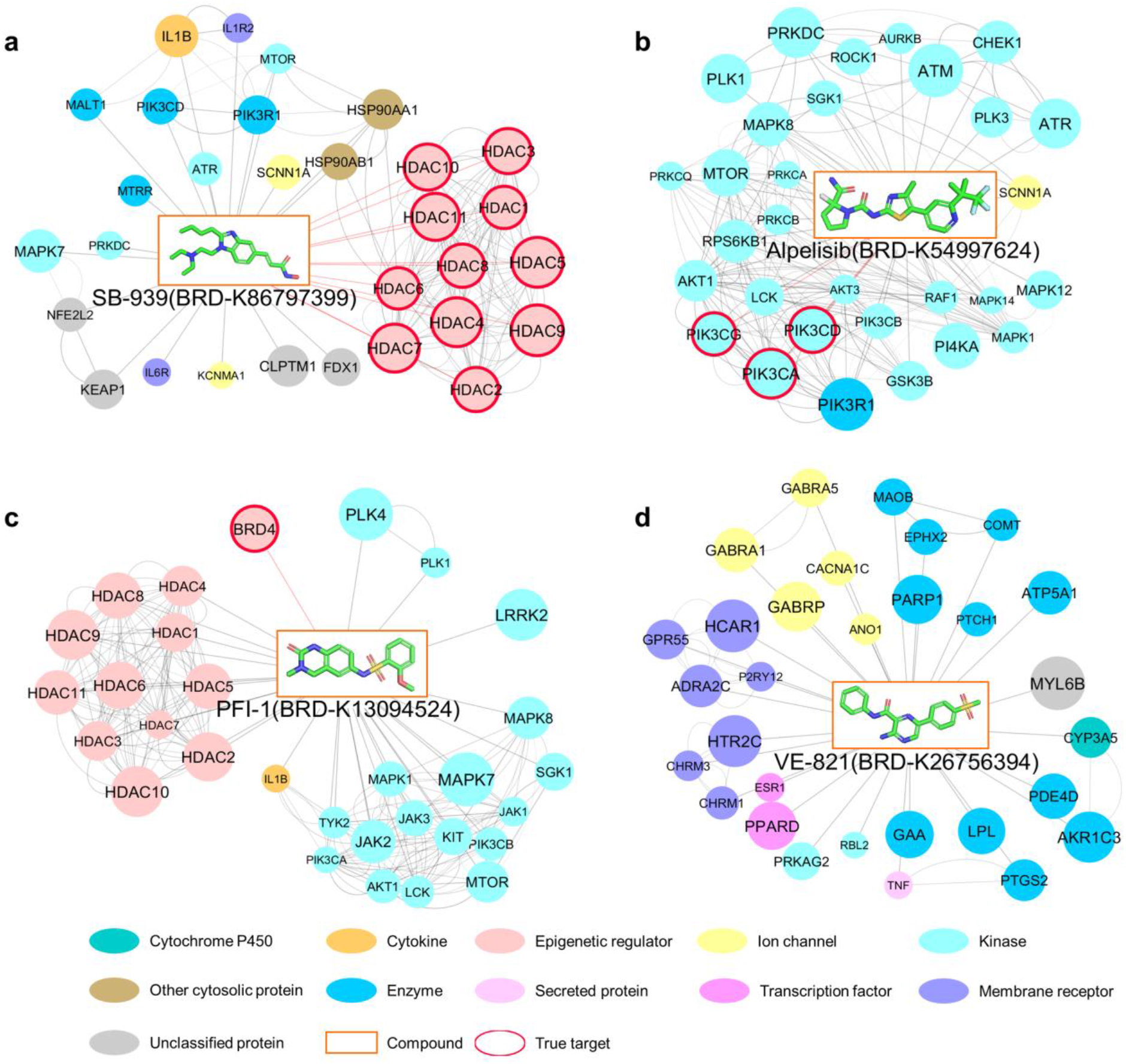
Examples of predicted targets (top 30) using the LINCS phase II data in PC3 cell lines. The following compounds were used for target prediction: (a) SB-939, (b) alpelisib, (c) PFI-1 and (d) VE-821. The nodes in rectangles represent compounds, and the nodes in circles represent the predicted targets. Predicted targets with a higher rank are indicated by a larger circle size. The links between predicted targets denote protein-protein interactions that are curated from the STING database with a combined score greater than or equal to 800.

## DISCUSSION

The drug-induced perturbation of cells leads to complex molecular responses upon target binding, such as the feedback loop that changes the expression level of the target node or its upstream and downstream nodes. These drug-induced responses likely resemble those produced after silencing the target protein-coding gene, which provides a rationale for comparing the similarity between chemical- and shRNA-induced gene expression profiles for target prediction (33). The encoding and denoising of a given experiment’s transcriptional consequences constitute a challenge. In this study, we proposed a new deep neural network model, the Siamese spectral-based graph convolutional network (SSGCN), to address this challenge.

The SSGCN model takes two differential gene expression networks (a chemical-induced network and a shRNA-induced network) as input and integrates heterogeneous experimental condition information to account for variances such as cell line-, dose- and time-dependent effects. By training on known compound-target interaction data, the model can automatically learn the hidden correlation between gene expression profiles, and this “deep” correlation was then used to query the reference library of 179,361 KD-perturbation profiles with the aim of identifying candidate target-coding genes. The pipeline achieved markedly improved target prediction performance on a benchmark test set. For more rigorous time-split validation using LINCS phase II data, the target prediction results obtained with our method were impressive compared with those achieved with the conventional CMap-based similarity approach. Furthermore, the pipeline only requires cell-level transcriptomic profiling data as input, which can be obtained in a high-throughput manner using commercial gene expression microarrays, RNA sequencing (RNA-Seq), and even the low-cost L1000 assay. Therefore, the methodology is not limited by the structural availability of proteins and is complementary to chemical similarity-based approaches. Overall, the SSGCN model allows *in silico* target inference based on transcriptional data and is of practical value for repositioning existing drugs or exploring not-well-characterized bioactive compounds and natural products.

## SUPPLEMENTARY DATA

Supplementary Data include: the numbers of KD genes in each KD time (Table S1), the predicted results in the PC3 cell line on the LINCS II data (Table S2) and a Venn diagram to show the target space of external test sets 1 and 2 (Figure S1).

## ACKNOWLEDGEMENTS

We gratefully acknowledge financial support from the National Natural Science Foundation of China (81773634 to M.Z), National Science & Technology Major Project “Key New Drug Creation and Manufacturing Program”, China (Number: 2018ZX09711002 to H.J.), and “Personalized Medicines—Molecular Signature-based Drug Discovery and Development”, Strategic Priority Research Program of the Chinese Academy of Sciences (XDA12050201 to M.Z.).

