## Supplementary Data for "Computational target fishing by mining transcriptional data using a novel Siamese spectral-based graph convolutional network"

**Table S1. The numbers of KD genes in each KD time**

| Cell lines | The number of genes | The number of genes (96h) | The number of genes (120h) | The number of genes (144h) |
|---|---|---|---|---|
| PC3 | 3980 | 3822 | 128 | 1725 |
| A549 | 3724 | 3724 | 0 | 0 |
| MCF7 | 3688 | 3471 | 0 | 1837 |
| HT29 | 3665 | 3665 | 0 | 0 |
| A375 | 3826 | 3826 | 0 | 0 |
| HA1E | 3801 | 3801 | 0 | 0 |
| VCAP | 4134 | 34 | 4121 | 0 |
| HCC515 | 3522 | 3522 | 0 | 0 |

**Table S2. The predicted results in the PC3 cell line on the LINCS II data**

| id | target | rank |
|---|---|---|
| A07563059 | ADRB2 | 48 |
| A12896037 | ADRA2C | 91 |
| A13021932 | YES1 | 77 |
| A13254067 | PPM1B;PPP1CC;PPP2CA;PTPN1;PPP2R5A | 584;1326;297;171;3335 |
| A16347691 | GMNN | 2219 |
| A28467416 | PIK3CB;MTOR;PIK3CA;PIK3CG;PIK3CD | 18;10;9;13;8 |
| A28545468 | EHMT2;MAOB | 14;67 |
| A29520968 | HSPB1 | 1770 |
| A48881734 | EZH2 | 1596 |
| A52922642 | CACNA1C | 201 |
| A64553394 | ADRB2 | 155 |
| A65730376 | DOT1L | 3764 |
| A82035391 | JUN | 378 |
| A82156122 | DPP4 | 771 |
| A82772293 | HRH1;HTR2C;CHRM3;CHRM1 | 2756;2354;2808;2367 |
| A86248581 | CDA | 1785 |
| A92800748 | TEK | 459 |
| A93093700 | LMNA | 1399 |
| K00152668 | RARB | 105 |
| K01577834 | ADORA2A | 525 |
| K01674964 | HRH1;BLM | 31;1314 |
| K02314383 | AR | 132 |
| K03194791 | PDE4D | 30 |
| K03390685 | MAP2K1 | 77 |
| K06762493 | GMNN;APEX1 | 1523;2360 |
| K07106112 | ERBB4;ERBB2;EGFR | 497;60;23 |
| K07310275 | AKT1;MTOR;PIK3CA | 13;12;1 |
| K07753030 | RGS4;BLM | 3736;3080 |
| K08109215 | BRD2;BRD3;BRD4 | 1413;2786;3 |
| K08248804 | XIAP | 88 |
| K08586861 | TBXA2R;MBNL1 | 297;3428 |
| K08832567 | GMNN;CA12 | 2544;50 |
| K08976401 | LMNA;NFKB1;APEX1;EHMT2 | 1322;341;3206;123 |
| K09372874 | IMPDH2 | 232 |
| K09711437 | PLA2G2A | 59 |
| K10859802 | GPR119 | 214 |
| K11267252 | RET;ALK | 395;760 |
| K12609457 | LMNA | 907 |
| K13094524 | BRD4 | 7 |
| K13662825 | CDK4;CDK9;CDK5;CDK1 | 34;58;13;18 |
| K14704277 | LMNA;BLM | 1697;1238 |
| K14870255 | AXL | 1696 |
| K15170068 | MAN2B1 | 1756 |
| K15179879 | HSPB1;AR;NFE2L2;PSMB8 | 2124;190;33;2208 |
| K15507868 | PPARG | 8 |
| K16295392 | EDNRB | 131 |
| K16730910 | EPHX2;BRAF | 182;24 |

| | | |
|---|---|---|
| K16761703 | PRKCE;PIM3;PKN2;CDK2;PRKCA;GSK3B;PIM2;FLT4;MAP4K4;RPS6KA3;PLK1;FER;PRKCB;PRKCZ;GSK3A;PRKCQ;PRKCH;STK3;MST1R;RPS6KB1;PIM1;CAMK2D;SLK;PRKCD;CAMK2G | 163;3858;197;16;89;7;3864;62;672;139;66;3764;101;236;21;155;438;407;2502;43;48;981;318;317;627 |
| K16803204 | TYK2;JAK1;FLT4;JAK3 | 879;1922;2217;493 |
| K17068645 | HDAC8;HDAC6;HDAC10;HDAC1;HDAC2;HDAC3 | 499;427;1035;469;1063;813 |
| K17203476 | FGFR1;KDR | 139;73 |
| K17498618 | LMNA | 1980 |
| K17555800 | MAPK14 | 7 |
| K18157228 | GMNN;NFE2L2 | 2951;129 |
| K18961567 | AXL | 2430 |
| K19412355 | LCK;HSPB1;RAF1;EGFR;CDK1;PRKCB;MAPK8;PRKCA;CSNK1D;MAPK14;MAPK9;MAP2K2 | 73;1291;133;81;23;303;165;250;2173;51;18;17 |
| K19601669 | TAOK2;MAP3K2;DAPK3;RPS6KA1;ULK1;ROCK1;PLK1;RPS6KA2;ULK2;PLK4;RPS6KA5;JAK3;TYK2;GAK;IRAK1;JAK2;CAMK2G;ROCK2;MKNK2;JAK1;DAPK1;CAMK2D;LRRK2 | 2820;490;2173;316;2791;284;814;111;3687;108;2213;33;373;323;1875;243;679;342;1885;327;627;588;193 |
| K19761926 | AR | 47 |
| K20079257 | CBX1 | 2700 |
| K20722021 | LCK;RAF1;EPHA2;ABL1;EGFR;RET;PDGFRB;KDR;BRAF;KIT | 37;117;203;129;27;168;139;60;257;120 |
| K20745393 | NFKB1 | 110 |
| K21025364 | MELK;IRAK4;INSR;IGF1R;MAPK8;ERBB4;DAPK3;CHEK2;CAMKK2;CDC7;ROS1;MAPK9;FES;MET;NEK4;ERBB2;PLK1;AURKA;PLK4;FER;CLK4;EGFR;MAPK1;ALK;PHKG2;PTK2;AURKB;CSNK2A1;CAMK2D;LRRK2 | 637;2891;56;119;16;92;3270;204;673;2186;232;130;3616;58;2138;49;3;93;91;3624;48;85;29;194;117;45;14;136;559;113 |
| K21295289 | KCNN4 | 410 |
| K21396683 | MMP2;MMP7;MMP14;MMP1;ADAM17 | 289;3229;296;3049;707 |

| | | |
|---|---|---|
| K21718444 | MELK;CSNK2A2;TTK;YES1;STK33;DYRK1A;PIP4K2B;STK10;MAP4K3;MAP3K2;NTRK1;DAPK3;PRPF4B;CHEK2;MAPK9;TBK1;MAP2K5;MYLK2;FLT3;ULK1;ABL1;MAP2K3;MET;ROCK1;CLK2;CDK7;RIPK1;MAP4K4;RET;RPS6KA3;SGK3;ACVR1;SRPK1;MAP2K4;PDGFRA;PLK4;ULK3;RIOK3;ULK2;STK11;CHUK;RIPK4;CLK4;KIT;STK17A;STK16;JAK3;TYK2;PHKG2;NEK6;MAP3K11;CLK1;GAK;NTRK3;KDR;AXL;ABL2;NEK7;FLT1;IRAK1;RPS6KB1;JAK2;AURKB;TNIK;ROCK2;PDGFRB;MARK4;MAP3K7;SLK;BMPR1B;LR | 799;89;617;100;2488;48;1751;318;1290;97;72;1721;653;149;40;118;463;481;12;2246;7;1655;39;46;1235;124;950;587;66;168;569;667;2262;245;1;198;3129;3336;3463;2474;205;996;26;9;81;3156;61;116;150;2690;745;2;201;239;65;3345;1910;3964;19;2054;75;30;49;200;141;32;1408;500;183;2007;76 |
| K21908111 | PIM2;PIM1;PIM3 | 3673;370;3841 |
| K22482860 | EHMT2 | 74 |
| K22822991 | CTSK;CTSH | 3530;2438 |
| K22861715 | CYP1A1;CYP1B1;CBR1 | 2772;161;1762 |
| K23190681 | YES1 | 195 |
| K23228615 | MAP3K9;PIK3CB;MTOR;PIK3CA;PIK3CG;PIK3CD;SYK | 2648;15;8;10;13;9;50 |
| K24723746 | CHRM3;CHRM1 | 3;5 |
| K25140590 | HPGD | 809 |
| K25361343 | MMP2;MMP7;MMP14;MMP1;ADAM17 | 1059;1745;582;2572;667 |
| K26603252 | ABL1;EGFR;MAPK1;ERBB2;ERBB4 | 208;19;153;26;227 |
| K26667523 | MAP2K1 | 9 |
| K26756394 | ATR | 1594 |
| K27182532 | MST1R;MET;KDR;AXL;FLT | 2162;135;73;3092;176 |
| K27204852 | CTSK;CTSD | 3892;370 |
| K27955832 | MGMT | 2157 |
| K28392481 | FGFR4;FGFR2;FGFR3;FGFR1;KDR | 1915;189;115;210;63 |
| K28965160 | MAPK1 | 102 |
| K29735307 | AR | 89 |
| K31135544 | ADORA2A;APEX1 | 39;1812 |
| K31309378 | FLT1;FGFR2;FLT4;FGFR1;KDR | 150;627;302;881;512 |
| K31313613 | CDK7 | 403 |
| K31812033 | ADRB2 | 125 |
| K31928526 | AURKB;RET;PDGFRB;PDGFRA;AURKA;KDR;KIT;FLT3 | 226;349;695;370;81;65;198;567 |
| K32285926 | MET | 232 |
| K32847255 | KDM6B | 1082 |
| K33251802 | GAA | 151 |
| K33610132 | EGFR | 10 |
| K35007173 | BLM | 2595 |
| K35245662 | AR | 49 |
| K35719256 | F3 | 1391 |
| K35775715 | HSPB1 | 1944 |
| K36280065 | JAK2;JAK3 | 96;71 |
| K36363294 | BRD2;BRD3;BRD4 | 2012;2919;16 |

| | | |
|---|---|---|
| K37111771 | GMNN | 3238 |
| K37764012 | PAK3;PAK1;PAK2;PAK4 | 3253;2246;39;2453 |
| K38852836 | HSP90AA1;TRAP1;HSP90AB1 | 2;578;1 |
| K38868394 | AR | 67 |
| K39252998 | ADRA2A;ADRA2C | 23;40 |
| K39974922 | FLT4;KDR;RET | 8;7;163 |
| K40718343 | FLT1;YES1;KDR;EGFR | 130;420;102;91 |
| K41599323 | HRAS | 2672 |
| K42436189 | ATR;MTOR | 2;9 |
| K42805893 | EGFR | 10 |
| K42828737 | LCK;RIOK2;TTK;IRAK4;YES1;STK33;DYRK1A;PIP4K2B;STK10;MAP4K3;MAP3K2;CHEK1;DAPK3;CSNK1G2;NTRK1;CSNK1D;PTK2B;CHEK2;CDK16;PRKAA1;MYLK;TBK1;CSNK1G3;MYLK2;MAP2K5;RPS6KA1;ULK1;FLT3;TAOK3;CLK2;STK25;FLT4;FGFR3;MAP4K4;RET;RPS6KA3;SGK3;SRPK1;RPS6KA2;PDGFRA;PLK4;ULK3;ULK2;MAP2K1;STK11;PRKAA2;CLK4;MAP2K2;RPS6KA5;MERTK;KIT;TYRO3;CSNK1A1;STK17A;STK16;CSNK1E;ALK;NUAK1;PHKG2;HIPK1;CLK1;STK39;GAK;KDR;AXL;FLT1;STK3;IRAK1;RPS6KB1;TNIK;PAK3;ROCK2;JAK1;CSNK2A1;PDGFRB;DAPK1;LYN;MAP3K7;SL | 40;1350;320;3207;75;1583;164;1325;136;778;211;37;3446;321;24;976;186;153;2845;555;1509;25;547;537;434;86;2466;15;3359;1248;3894;2;23;792;28;279;478;1103;837;9;48;3158;3766;46;2218;1362;67;57;1140;3316;3;2288;449;91;3426;480;238;2761;112;2523;152;538;105;53;3324;7;128;1698;30;525;2587;66;65;117;27;389;59;429;88;5;210;54 |
| K42948882 | CXCR4 | 2248 |
| K43002773 | AKT3;AKT2;AKT1 | 22;42;20 |
| K43802723 | FFAR1 | 187 |
| K43966364 | LMNA | 994 |
| K44309363 | ABCB1 | 70 |
| K44408410 | RGS4;BLM | 1202;1197 |
| K44827188 | SMO;SHH | 610;2888 |
| K44844162 | PIK3CB;PIK3CA;PIK3CG;PIK3CD;PIK3C2B | 14;10;12;8;1511 |
| K45275534 | F10 | 62 |
| K45924332 | GMNN;LMNA | 1828;1123 |
| K46290096 | ADORA2A | 69 |
| K46692793 | CHRM3;CHRM1 | 94;87 |
| K47642186 | PDE4D | 9 |
| K48068743 | DPP4 | 356 |
| K48213016 | MAPK9;MAPK8 | 2731;903 |
| K48461310 | GMNN;LMNA | 1722;1280 |
| K48894757 | TBXA2R | 120 |
| K49055432 | PIK3CA | 280 |
| K49215523 | PIK3CB | 493 |

| | | |
|---|---|---|
| K50000283 | MAPKAPK2;CDK1;CDK9;DYRK1A;CSNK1D;PKN2;CSNK1G2;DAPK3;CDC7;CDK2;GSK3B;CSNK1G3;ROCK1;CLK2;CLK4;GSK3A;CDK8;CSNK1A1;STK17A;MAPK12;CDK5;PRKCQ;DYRK3;IRAK1;ROCK2;MKNK2;MA | 78;14;40;81;1900;215;260;2771;1569;10;48;216;103;1151;87;94;2478;1298;114;49;8;89;1580;2093;166;238;605 |
| K51263939 | GMNN | 1303 |
| K51544265 | MET;RET;TEK;KDR;AXL;KIT;FLT3 | 47;89;308;46;2782;77;53 |
| K51747290 | EHMT2 | 445 |
| K52183142 | AR | 82 |
| K52233191 | GSK3A;TAOK3;CDK5;CDK1;CLK2;CASK;CDK7;CDK4;CDK9;CLK1;DYRK1A;MAPK7;SLK;CLK3;CDK16;CDK2;CLK4;GSK3B | 115;2932;9;16;955;710;199;46;114;3;66;1126;351;3023;3097;6;52;39 |
| K52818472 | CFTR | 418 |
| K53417444 | MELK | 1150 |
| K53508936 | MMP2;MMP7;MMP14;MMP1;ADAM17 | 53;1342;55;1438;301 |
| K53581288 | TYK2;JAK1;JAK2;JAK3 | 263;237;372;232 |
| K53814070 | ROCK2;ROCK1 | 744;352 |
| K53963539 | HRH1;HTR2C | 21;1 |
| K54520417 | NR1H3;NR1H2 | 145;169 |
| K54606188 | BRD2;BRD3;BRD4 | 2163;2569;17 |
| K54640016 | NR1H4 | 315 |
| K54997624 | PIK3CG;PIK3CD;PIK3CA | 10;7;3 |
| K55013654 | HTR2C | 1 |
| K55512740 | MST1R;MET | 397;277 |
| K56032964 | INSR;ROS1;IGF1R;ALK | 41;306;113;283 |
| K56195681 | MAPK14 | 17 |
| K56405753 | FLT1;MST1R;MET;FGFR2;FLT4;JAK2;FGFR3;FLT3;FGFR1;NTRK1;PDGFRA;AURKA;KDR;MERTK | 53;659;124;97;5;17;81;67;14;60;8;41;32;3033 |
| K57169635 | ERBB4;ERBB2;EGFR | 230;49;15 |
| K57754230 | EZH2 | 1536 |
| K58114536 | MAOB | 63 |
| K58435339 | HSP90AB1 | 1 |
| K58486055 | CXCR2 | 1279 |
| K58501140 | FFAR1 | 88 |
| K59325863 | PSMB2;PSMB5 | 3;4 |
| K59433843 | TBXA2R | 73 |
| K59506194 | GMNN | 2424 |
| K59573506 | AR;NR3C1 | 64;93 |
| K60997853 | LCK;CDK5;CDK7;CDK4;NTRK1;CDK2 | 2;13;230;104;51;35 |
| K61195623 | ABCB1;TUBB3 | 274;109 |
| K61397605 | HDAC2;HDAC1 | 364;419 |
| K62391742 | BCL2;BCL2L1 | 44;17 |
| K62627508 | MDM2 | 213 |
| K62762455 | AR | 224 |
| K63126190 | ADRA2A;ADRB2 | 39;21 |
| K63861289 | BLM;HIF1A | 2547;234 |
| K64538373 | MAP2K1 | 183 |
| K64925568 | MDM2 | 156 |

| | | |
|---|---|---|
| K65182930 | HSP90AA1;HSP90B1;TRAP1;HSP90AB1 | 2;6;505;1 |
| K67121414 | F10 | 5 |
| K67174965 | MAPK1 | 71 |
| K67525671 | APEX1 | 1868 |
| K68232650 | SCN9A | 257 |
| K68532323 | KIF11 | 8 |
| K68938568 | XIAP | 34 |
| K69001009 | KDR;MET | 38;147 |
| K69776681 | BRD4;PLK1 | 2;19 |
| K70301465 | BTK;FGFR2;EGFR;YES1;RET;LYN;FLT3 | 138;248;20;395;442;653;418 |
| K71075609 | CA2;LMNA | 40;628 |
| K71221037 | BIRC2 | 3387 |
| K71281111 | HSP90AA1;HSP90AB1 | 2;1 |
| K71480163 | AKT3;AKT2;AKT1 | 11;26;22 |
| K71822263 | BLM | 2785 |
| K73107279 | BLM | 286 |
| K73237276 | VDR | 43 |
| K73381542 | HIF1A;TP53;IDH1;GAA | 306;139;814;62 |
| K74339692 | ADRA2A;ADRA2C | 59;27 |
| K74363950 | CHRM3;CHRM1 | 17;45 |
| K75844781 | HTR2C;BLM | 86;3397 |
| K76401790 | MAPK9 | 1045 |
| K76694128 | YES1 | 97 |
| K76841105 | RARA;RARG;RARB | 171;303;233 |
| K77396579 | DPP4 | 68 |
| K77625799 | LCK;ERBB3;IRAK4;YES1;STK10;MAP2K5;ABL1;FLT4;MAP4K4;RET;RIPK2;ACVR1;FGFR1;PDGFRA;EPHA6;KIT;TYRO3;EGFR;GAK;KDR;AXL;ABL2;FLT1;ROCK2;PDGFRB;LYN;SLK;BMPR1B; | 35;3542;3492;111;175;414;63;20;492;90;123;1698;77;25;2944;66;1128;52;118;47;2688;1800;76;293;36;199;285;1966;78;50 |
| K78055238 | GSK3A;LCK;CDK2;GSK3B | 34;9;11;14 |
| K78118466 | TGFBR1;RIPK2;ACVR1B | 155;344;1380 |
| K78790801 | NTRK1;AXL;KIT;FLT3 | 216;1337;248;331 |
| K79018442 | CHUK;IKBKB | 669;344 |
| K79102359 | HTR2C | 8 |
| K79239947 | CDK7;PTK2 | 47;334 |
| K80082640 | F10 | 149 |
| K80343549 | ERBB4;ERBB2;EGFR | 253;52;26 |
| K80608265 | ESR2;ESR1 | 161;98 |
| K80700417 | FLT3;PIM1;PIM3 | 94;380;3440 |
| K81458380 | PIM2;CSNK2A2;PIM1;CSNK2A1;DAPK3;CLK3;TBK1;FLT3 | 3735;98;149;182;3121;2430;83;51 |
| K81672972 | PTGFR;TBXA2R | 24;141 |
| K82603084 | EHMT2 | 102 |
| K82928847 | HDAC8;HDAC6;HDAC10;HDAC1;HDAC2;HDAC3 | 9;11;6;10;8;7 |
| K83699324 | PLK1 | 8 |
| K84564571 | FLT4;KDR | 149;48 |
| K84794093 | GMNN | 735 |
| K86118762 | INSR;IGF1R | 300;121 |
| K86525559 | YES1;CHEK1;CHEK2 | 55;10;108 |

| | | |
|---|---|---|
| K86797399 | HDAC8;HDAC7;HDAC4;HDAC6;HDAC10;HDAC1;HDAC2;HDAC9;HDAC5;HDAC11;HDAC3 | 9;4;3;11;6;10;8;1;2;5;7 |
| K86816506 | FAAH | 5 |
| K86856088 | EHMT2 | 153 |
| K87465484 | EHMT2 | 242 |
| K87782578 | BTK | 637 |
| K87904882 | PRKCE | 690 |
| K88025533 | S1PR3 | 684 |
| K88052444 | PPARD | 134 |
| K88506063 | ROCK2;CSNK2A1 | 394;163 |
| K88807631 | ABCB1;ABCG2 | 127;45 |
| K89208535 | ADRB2 | 47 |
| K89561498 | ABCG2;TOP1 | 206;1 |
| K90195324 | ESR2;ESR1 | 22;19 |
| K90777969 | EBP | 518 |
| K91308639 | AR | 55 |
| K92213669 | MTTP | 1829 |
| K92441787 | RARA;RXRG;RXRA;RXRB;AR;RARB | 90;6;218;12;51;16 |
| K92613113 | MAOB | 84 |
| K93123848 | RAF1;TAOK2;CIT;RET;PDGFRB;STK10;BRAF;KIT;DDR | 234;3915;1364;323;203;337;136;166;255 |
| K93779381 | PRKCD | 8 |
| K94534639 | BACE1 | 66 |
| K95880107 | MAPK14 | 135 |
| K96550715 | CHRM1;DPP4 | 25;723 |
| K96936751 | APEX1 | 475 |
| K97746869 | LMNA | 1307 |
| K97963946 | MET | 135 |
| K98004573 | HRH1 | 34 |
| K98453471 | LMNA;NFKB1;EHMT2 | 1186;36;40 |
| K98572433 | ERBB2;ERBB3;EGFR | 85;2856;140 |
| K99023089 | AKT3;ROCK2;AKT2;AKT1 | 10;122;28;21 |
| K99092662 | AR | 16 |
| K99113996 | MTOR | 9 |
| K99475619 | TOP1 | 1 |
| U07805514 | LCK;ABL1;YES1;SRC;KIT | 10;40;200;137;58 |
| U70626184 | RPS6KA4;MYLK;PLK2;PIP4K2C;PLK1;ALK;DAPK3;CAMKK2;GAK;PTK2B;PLK3;BRD4;RPS6KA5;PTK2 | 3480;1508;167;967;39;220;3545;843;82;189;79;128;1141;63 |

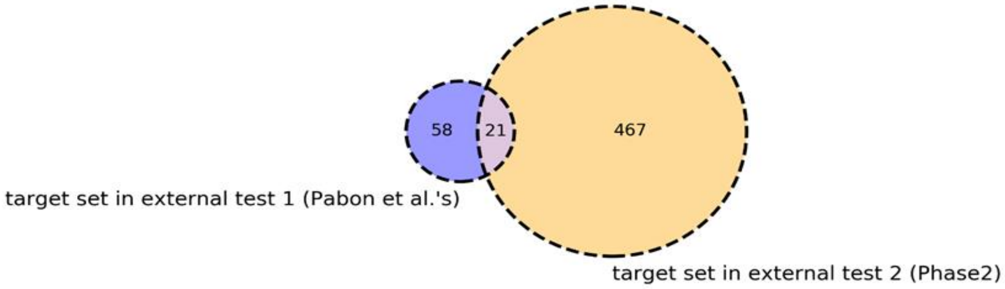


**Figure S1. Venn diagram to show the target space of external test sets 1 and 2.**
